# Termite mounds contain distinct methanotroph communities that are kinetically adapted to elevated methane concentrations

**DOI:** 10.1101/717561

**Authors:** Eleonora Chiri, Chris Greening, Stefan K. Arndt, Philipp A. Nauer

**Affiliations:** School of Ecosystem and Forest Sciences, University of Melbourne, Richmond, VIC 3121, Australia; School of Biological Sciences, Monash University, Clayton, VIC 3800, Australia; School of Chemistry, Monash University, Clayton VIC 3800, Australia

## Abstract

Termite mounds have recently been confirmed to mitigate approximately half of termite methane (CH_4_) emissions, but the aerobic methane-oxidizing bacteria (methanotrophs) responsible for this consumption have not been resolved. Here we describe the abundance, composition, and kinetics of the methanotroph communities in the mounds of three distinct termite species. We show that methanotrophs are rare members of the termite mound biosphere and have a comparable abundance, but distinct composition, to those of adjoining soil samples. Across all mounds, the most abundant and prevalent particulate methane monooxygenase sequences detected were affiliated with Upland Soil Cluster α (USCα), with sequences homologous to *Methylocystis* and Tropical Upland Soil Cluster also detected. The Michaelis-Menten kinetics of CH_4_ oxidation in mounds were estimated from *in situ* reaction rates. The apparent CH_4_ affinities of the communities were in the low micromolar range, which is one to two orders of magnitude higher than those of upland soils, but significantly lower than those measured in soils with a large CH_4_ source such as landfill-cover soils. The rate constant of CH_4_ oxidation, as well as the porosity of the mound material, were significantly positively correlated with the abundance of methanotroph communities of termite mounds. We conclude that termite-derived CH_4_ emissions have selected for unique methanotroph communities that are kinetically adapted to elevated CH_4_ concentrations. However, factors other than substrate concentration appear to limit methanotroph abundance and hence these bacteria only partially mitigate termite-derived CH_4_ emissions. Our results also highlight the predominant role of USCα in an environment with elevated CH_4_ concentrations and suggest a higher functional diversity within this group than previously recognised.

## Introduction

Termites are mound-building eusocial insects that live in colonies throughout the tropics and subtropics. These organisms completely degrade lignocellulose in a process primarily mediated by anaerobic symbiotic microorganisms in their hindgut [1]. During this process, hydrogenotrophic methanogens produce substantial amounts of methane (CH_4_) that is emitted from the termite into the atmosphere [2–4]. Production rates vary by three to four orders of magnitude depending on the termite species and their dietary preferences (i.e. wood-, grass-, soil- or fungus-feeding) [1, 4, 5]. Current models suggest that termites are responsible for between 1 to 3 % of global CH_4_ emissions to the atmosphere [6].

Aerobic methane-oxidizing bacteria (methanotrophs) significantly mitigate emissions of CH_4_ from termites [7]. Methanotrophs gain carbon and energy by oxidising CH_4_ to carbon dioxide, with the first step in this reaction being catalysed by particulate and soluble methane-monooxygenases [8]. It is controversial whether termite hindguts harbor such organisms; while *Methylocystis* spp. were recently isolated from termites [9], other studies could not detect methanotroph functional gene markers or measurable amounts of ^14^CO_2_ during ^14^CH_4_ incubation experiments [10]. We also observed that the addition of inhibitors of CH_4_ oxidation did not increase direct termite CH_4_ emissions [7]. However, many termite colonies construct large mounds built from soil material or build their nest in soil, which is generally a sink for atmospheric CH_4_ [11]. While results from incubation experiments of mound material were conflicting [12–14], we recently presented clear evidence of widespread CH_4_ oxidation in North Australian termite mounds [7]: results from three different *in situ* methods to measure CH_4_ oxidation in mounds confirmed that methanotrophs mitigate between 20 to 80 % of termite-derived CH_4_ before emission to the atmosphere. However, the community composition and kinetic behaviour of the methanotrophs responsible remain largely unknown.

Compared to soils, methanotrophs inhabiting termite mounds have received little attention. Ho et al. [14] investigated mound material of the African fungus-feeding termite *Macrotermes falciger* using a *pmoA*-based diagnostic microarray approach. Community composition differed between mound and soil, and at some locations within the large compartmentalised mounds. Slurry incubations confirmed potential CH_4_ oxidation at high and low CH_4_ concentrations. The mound community was reportedly dominated by gammaproteobacterial methanotrophs of the JR3 cluster, while the functional gene of the soluble methane-monooxygenase could not be detected, nor could methanotrophs of the Verrucomicrobia and Methylomirabilota (NC10) phyla. Beside this pioneering work, the mound methanotroph communities of no other termite species has been investigated. However, large differences might exist between termite species and particularly their dietary preferences, as well as the mounds’ impressive variety of sizes, shapes and internal structures [15, 16]. Reflecting this, *in situ* studies have shown that there is a large variation in methanotroph activity in mounds both within and between species; for example, *Tumulitermes pastinator* mounds appear to be largely inactive and still a high fraction of termite-derived CH_4_ can be oxidised in soil beneath mounds, due to facilitation of CH_4_ transport within the mound [7]. It remains unclear whether differences in methanotroph community abundance or composition account for these activity differences.

In this work, we aimed to resolve these discrepancies by conducting a comprehensive analysis of the composition and kinetics of the methanotroph communities within termite mounds of Australian termite species. Three mound-building termite species were selected, the wood-feeding *Microcerotermes nervosus* (Mn), soil-interface-feeding *Macrognathotermes sunteri* (Ms), and grass-feeding *Tumulitermes pastinator* (Tp), which represent the three main feeding groups present in Australia [17]. Mounds of these species were previously confirmed to oxidise a high fraction of termite-produced CH_4_ [7]. We used the *pmoA* gene, encoding a subunit of the particulate methane monooxygenase present in most methanotrophs [18], as a molecular marker to study the abundance, diversity, and composition of the methanotrophs within 17 mounds and a subset of adjoining soils. In parallel, we performed *in situ* studies using gas push-pull tests (GPPTs) to derive the kinetic parameters of CH_4_ oxidation. We demonstrate that methanotrophic communities in the core and periphery of termite mounds are compositionally and kinetically distinct from those of surrounding soil, and primarily comprise methanotrophs affiliated with the Upland Soil Cluster α (USCα) with an apparent medium affinity for CH_4_.

## Materials and Methods

### Field sites and sampling

Field tests and sampling were performed in April and May 2016 in a coastal savanna woodland on the campus of Charles Darwin University in Darwin, Northern Territory, Australia (12.370° S, 130.867° E). The site is described in detail in Nauer et al. [7]. For this study, 29 mounds were first subject to *in situ* methane kinetic measurements using gas push-pull tests (described below). For further investigations following field measurements, we selected 17 termite mounds of an appropriate size for processing in the laboratory (initially 18, but one was damaged during transport and had to be discarded). These mounds were first excavated but kept intact to measure internal structure, volume, densities, and porosities as previously described [19]. They were then deconstructed to (i) sample termites for species identification, (ii) collect mound material for gravimetric water content measurements, and (iii) collect mound material for molecular analyses of methanotrophic community. For species determination, soldiers were individually picked and stored in pure ethanol for species confirmation as previously described [19]. For gravimetric water content measurements, approximately 200 g of mound material from both core and periphery locations were subsampled and oven dried at 105 °C for >72 hours; subsamples were measured before and after drying and the water content calculated based on mass loss. Subsamples for physicochemical parameters were oven-dried at 60 °C for 72 h, carefully homogenised into a composite sample for each termite species and location, and sent to an external laboratory for analyses according to standard protocols (CSBP laboratories, Bibra Lake WA, Australia). For community analysis, mound and soil material was collected under sterile conditions using bleach- and heat-sterilised spatulas, and immediately stored in autoclaved 2 mL centrifuge tubes at −20 °C. For each sampling location (mound core and periphery, soil beneath and surrounding the mound), we collected triplicates of pooled materials deriving from three different spots. Mound cores were sampled from within 20 to 30 cm from the approximate centroid of mound, whereas mound periphery was collected from the outer 5 to 10 cm of the mound. For a subset of the investigated mounds, soil was collected from beneath the mound immediately after mound excavation, and from the surrounding soil within a 1-2 m radius from the mound.

### Genomic DNA isolation and *pmoA* gene amplification

Each individual sample of mound material and soil was homogenised. DNA was extracted from 0.25 to 0.5 grams of each sample using the PowerLyzer PowerSoil DNA Isolation Kit (Qiagen, US), according to the manufacturer instructions. The purity and integrity of the DNA extracts was verified by spectrophotometry (NanoDrop ND-1000 spectrophotometer, Nanodrop Technologies Inc., US) and PCR amplification of 16S rRNA genes. Good yields of high-quality, amplifiable genomic DNA were obtained from all 48 samples. Amplicons of the *pmoA* gene were also obtained from all samples using previously described degenerate primers (A189f 5′-GGNGACTGGGACTTCTGG-3′, and mb661 5′-CCGGMGCAACGTCYTTACC-3′) [20, 21] and cycling conditions [22]. Amplification reaction mixtures (25-50 µL final volume) were prepared using 1 µl of DNA extract as template, 1 × PCR buffer, 0.2 mM of each primer, 0.25 mM deoxynucleoside triphosphates (dNTPs), and 0.025 U µl^−1^ of Taq polymerase (Takara Biotechnology Ltd., Japan). Different dilutions (undiluted to 1:100 in PCR-grade water) of DNA extracts were used as template during amplification, and the dilution resulting in the highest yield and quality of PCR product was used for further analyses.

### Quantitative PCR assays

Quantitative PCR assays were used used to estimate the abundance of the total bacterial community and methanotroph community. Total bacterial abundance was estimated by amplifying the 16S rRNA gene using degenerate primers (515FB 5’-GTGYCAGCMGCCGCGGTAA-3’ and 806RB 5’-GGACTACNVGGGTWTCTAAT-3’) and cycling conditions as previously described [23–25]. Methanotroph abundance was estimated by amplifying the *pmoA* gene using previously described degenerate primers (A189f 5′-GGNGACTGGGACTTCTGG-3′, and mb661 5′-CCGGMGCAACGTCYTTACC-3′) [20, 21] and cycling conditions [22]. Gene copy numbers were determined using a LightCycler 480 real-time PCR system (Roche, Basel, CH). Individual reactions contained 1 × PowerUp SYBR Green Master Mix (Thermo Fisher Scientific), 400 µM of each primer, and 1 µl of diluted environmental DNA mixed to a final volume of 20 µl. Thermal profiles were adapted from those used for previous PCRs and included an acquisition step of 85 °C for 30 s at the end of each amplification cycle. Melting curve analysis was performed as follows: 95 °C for 15 s, 60 °C for 60 s, 95 °C for 30 s, 60 °C for 15 s. For each assay (96-well plate), duplicate serial dilutions of quantified DNA extract from *Methylosinus trichosporium* were used for calibration curves to quantify 16 rRNA genes or *pmoA* genes. Each sample was analyzed in triplicate, and a total of three assays were required for each gene to include all the samples. Amplification efficiencies calculated from the slopes of calibration curves were >70% and R^2^ values were >0.98.

### Methanotroph community analysis

The structure of the methanotroph community within each sample was inferred from amplicon sequencing of community *pmoA* genes. DNA extracts of all samples were sent to the Australian Genome Research Facility (AGRF; Brisbane, QLD) for preparation of *pmoA* gene amplicon libraries using the above primer sets (A189f and mb661) and thermal conditions. Subsequent amplicon sequencing was performed on a MiSeq DNA-sequencing platform using a 600-cycle MiSeq Reagent Kit v3 (Illumina, San Diego, CA). Sequencing yielded 5,594,739 paired end sequences, of which 3,078,335 passed quality checks and data processing, and were used for subsequent analyses. Sequence read-counts spanned three orders of magnitude (10^5^ to 10^2^), with 54% of the samples exhibiting read counts above the average read-count value (>65K) and most mound periphery samples having read-counts below 10K. Six samples with read-counts below 1K and were excluded from subsequent analyses. Sequencing data was processed according to our previously published pipeline [26], with minor modifications. Briefly, reads were 3’-trimmed to remove ambiguous or low-quality endings, then merged and primer-site trimmed. Quality filters included an amplicon-size selection (471 nt) and the removal of amplicons containing stop codons (e.g., TAA, TAG, TGA). Sequences were also checked for correct open reading frames (ORF) using the FrameBot tool (http://fungene.cme.msu.edu/FunGenePipeline/framebot/form.spr). The centroid clustering method [27] identified 25 Operational Taxonomic Units (OTUs) that shared 86% nucleotide sequence similarity [28] with sequences from a curated *pmoA* gene database derived from Dumont et al., 2014 [29]. Phylogenetic distances of the assigned OTUs in relation to reference *pmoA* sequences were assessed as previously described [26]. The phylogenetic tree of protein-derived *pmoA* sequences was constructed using the maximum-likelihood method and the LG empirical amino acid substitution model, which showed the lowest Akaike information criterion (AIC) during substitution model testing, and was bootstrapped using 100 bootstrap replicates. All sequences affiliated with methanotrophs and no *pxmA* or *amoA* sequences were detected.

### Phylogenetic and diversity analyses

Alpha and beta diversity calculations, as well as read count normalization of the *pmoA* sequences, were performed with the package phyloseq v1.12.2 [30] from the open source software Bioconductor. To account for differences in numbers of reads between samples, we rarefied OTU counts to an even sampling depth of 1,611 read counts. Chao1, Shannon, and Inverse Simpson indices were computed to assess the alpha diversity of MOB communities. Beta diversity of methanotroph communities was measured using the phylogenetic metric Unifrac weighted by the relative abundance of individual OTUs (Lozupone and Knight, 2005). Differences were visualised using non-parametric multi-dimensional scaling ordinations (nMDS). To determine whether the observed between-group distances were statistically significant, we performed permutational multivariate analysis of variance (PERMANOVA) with the software PRIMER-E v7 (PRIMER-E Ltd., Plymouth, United Kingdom). Negative binomial models were performed on the non-rarefied OTU dataset to assess the differential abundance of bacterial OTUs between sample groups, and the false discovery rate approach was used to account for multiple testing.

### Gas push-pull tests

The gas push-pull test was used to estimate *in situ* activity coefficients as described previously [19, 31]. Michaelis-Menten parameters estimated from *in situ* methods are integrated measures across a large mass of substrate and are thus better suited to characterise the kinetic potential of whole microbial communities in heterogeneous systems than laboratory microcosms, which suffer from inevitable sampling bias [32]. In brief, a gas mixture containing laboratory air, ∼900 µL L^−1^ of CH_4_, and ∼0.1 L L^−1^ argon (Ar) was injected at a rate of ∼0.5 L min^−1^ into the lower center of the termite mounds and then immediately extracted from the same location at the same flow rate. During extraction, the injected gas mixture was gradually diluted with termite-mound air down to background levels; the tracer Ar accounted for this dilution due to its similar transport behavior to CH_4_. A timeseries of CH_4_ and Ar concentrations was collected during the 24 min injection phase, and the 36 min extraction phase. Concentrations of CH_4_ were measured quasi-continuously (frequency of 1 Hz) using a field-portable spectrometer (Fast Greenhouse Gas Analyser Los Gatos Research, Mountain View, CA). For Ar, discrete samples were collected at fixed intervals during injection (n = 3) and extraction (n = 10-12), as well as prior to injection to determine background levels. Argon concentrations were analysed on a customised gas chromatography system (SRI 8610, SRI Instruments, Torrance, CA) with an external thermal conductivity detector (TCD; VICI Valco Instruments Co., Houston, TX). To improve separation of Ar, oxygen was removed from the sample-gas stream prior to separation with a manually packed Pd-Al catalyst column [19, 33].

### Kinetic analysis

First-order rate coefficients of CH_4_ oxidation (activity coefficient *k*) were estimated from GPPTs via the slope of the logarithm of relative CH_4_ vs Ar concentrations, plotted against a transformed reaction time, according to the plug-flow reactor model for simplified GPPT analysis [34]. This data also allowed the calculation of reaction rates at different CH_4_ concentrations from segments of the extraction time-series, and thus the estimation of Michaelis-Menten parameters [35]. The running average of CH_4_ concentrations *C*_*CH4*_ over three consecutive extraction samples was multiplied with the corresponding activity (*k*) from the logarithmic plot to calculate an individual reaction rate *R*_*ox*_ for each segment. The Michaelis-Menten parameters *K*_m_ and *V*_max_ were then estimated for each individual GPPT, and for all combined pairs of concentrations and reaction rates, by fitting the Michaelis-Menten model *R*_*ox*_ = *V*_max_ * *C*_*CH4*_ / (*K*_m_ + *C*_*CH4*_) to the data using the non-linear regression routine nls() in R [36]. The AIC was calculated and compared with a linear regression model of *R*_*ox*_ vs *C*_*CH4*_; if the AIC of the linear model was lower, no Michaelis-Menten parameters were reported. Cell-specific activities and rates were calculated from reaction rates based on total mound dry mass, divided by *pmoA* copy numbers and assuming two gene copies of *pmoA* per cell. Correlations between kinetic parameters (*k, K*_*m*_, *V*_*max*_), gene abundance (*pmoA* and16S), and physical mound parameters (mound porosities, volume, water content) were tested for significance using linear regression, after transformations (sqrt for kinetic parameters, log for gene abundances) and removal of outliers as indicated by diagnostic plots (qq- and Cook’s distance).

## Results and Discussion

### Methanotrophic bacteria are in low abundance in termite mounds and associated soils

Quantitative PCR was used to estimate the abundance of the methanotroph community (*pmoA* copy number) and total bacterial community (16S gene copy number) in each mound and soil sample. Bacterial abundance was consistently high (av. 2.7 × 10^10^ 16S copy numbers per gram of dry soil; range 2.5 × 10^8^ to 3.4 × 10^11^) and did not significantly differ between sample locations **(Fig. 1b & Fig. S1)**; an earlier study found higher microbial biomass in the mound compared to soil [37], but this may reflect different methodologies applied to each substrate. In contrast, *pmoA* copy number was relatively low across the samples (av. 1.5 × 10^6^ copies per gram of dry sample material; range: 2.0 × 10^4^ to 1.8 × 10^7^) and just 0.0076% that of 16S copy number (range: 0.00018% to 0.048%) **(Figure 1a)**. Such values are comparable to those previously reported for the abundance of *pmoA* genes in upland soils that mediate atmospheric CH_4_ oxidation (∼10^6^ copy number, ∼0.01% relative abundance [38]).

**Figure 1.**
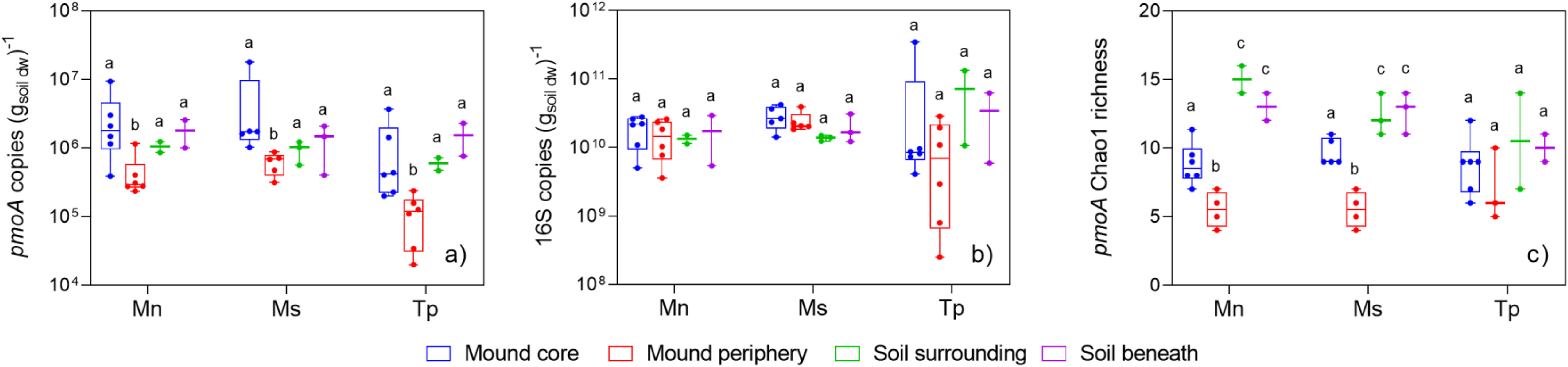
Abundance and richness of methane-oxidising bacteria (methanotrophs) in the mounds and adjoining soils. a) Abundance of the methanotroph community, based on copy number of the *pmoA* gene encoding the particulate methane monooxygenase 27 kDa subunit gene as determined by qPCR. b) Abundance of the total bacterial and archaeal community, based on copy number of the universal 16S rRNA gene (V4 region) as determined by qPCR. c) Estimated richness of the methanotroph community, based on Chao1 index of the *pmoA* gene as determined by amplicon sequencing. Mound samples (core and periphery) and adjoining soil samples (surrounding and beneath) were tested from mounds of three different termite species (*Microcerotermes nervosus*, Mn; *Macrognathotermes sunteri*, Ms; *Tumulitermes pastinator*, Tp). The box plots show minimum, lower quartile, median, upper quartile, and maximum values, with all individual values shown. Different letters denote significant differences between sample groups (*p* < 0.05, Wilcoxon signed-rank test).

Some differences in methanotroph abundance were observed between sample locations and termite species. Overall, *pmoA* copy number was 3.5-fold higher in mound core and 1.5-fold higher in soil beneath than in surrounding soil, though differences were below the threshold of significance **(Fig. S1)**. In contrast, *pmoA* copy numbers were significantly lower in mound periphery samples of all species (*p* = 0.028) **(Fig. S1)** and in mound samples of *T. pastinator* compared to the other two species tested (*p* = 0.028) **(Fig. S2)**; the latter observation is in line with the finding that CH_4_ oxidation occurs at low rates in *T. pastinator* mound material [7]. Consistently, *pmoA* copy numbers were significantly positively correlated with the porosity of the mound material, with periphery samples showing a stronger dependence (R^2^ = 0.49, *p* = 0.0051) than core (R^2^ = 0.26, *p* = 0.035); this suggests that denser mound material, as found in mound periphery and *T. pastinator* mounds, limits methanotroph abundance [19]. These differences may also reflect the relatively harsh conditions in the mound periphery, with its strong fluctuations of temperature and water content, compared to the core with termite-engineered homeostasis [39, 40]. Other physical parameters did not correlate with *pmoA* or 16S copy numbers.

### Termite mound methanotroph communities are compositionally distinct from those of associated soils

The composition and diversity of the methanotroph community in each sample was inferred through amplicon sequencing of the *pmoA* gene. Across the samples, 25 OTUs were detected **(Figure 2)**. Observed and estimated richness of these OTUs was higher in soil samples compared to mound samples (av. Chao1 of 9.0 for mound core, 5.9 for mound periphery, 12.3 for soil samples; *p* < 0.001) **(Figure 1c)**; however, these differences were driven primarily by rare OTUs in soil samples, with Shannon and inverse Simpson indices similar between samples **(Figure S3)**. Beta diversity of the samples was analysed by weighted Unifrac and visualised on an nMDS ordination plot **(Figure 3a)**. PERMANOVA analysis confirmed communities significantly differed between sample locations (*p* = 0.001) and termite species (*p* = 0.022). With respect to sample location, communities within mound core and periphery samples were similar and were compositionally distinct from soil communities; in addition, methanotroph communities in soils beneath mounds were more similar to those within mounds than those in soils surrounding mounds. This confirms previous inferences that mound and soil communities are different and shaped by termite activity [14]. In addition, core and peripheral mound communities significantly clustered by termite species, while soil samples did not; mound communities of *M. nervosus* and *T. pastinator* were more closely related than those of *M. sunteri* **(Figure 3a)**.

**Figure 2.**
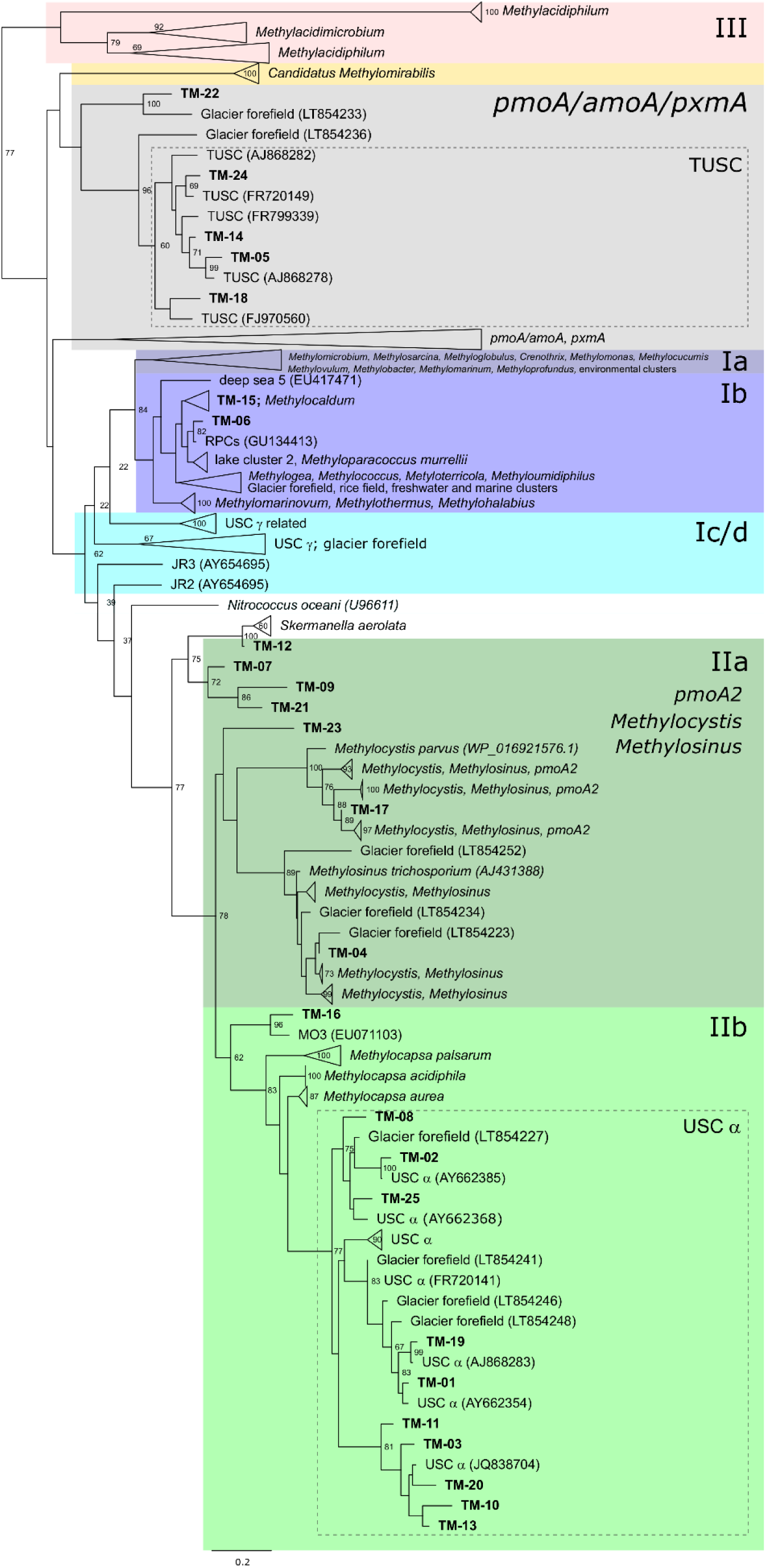
Maximum-likelihood tree showing the phylogenetic affiliation of the protein-derived *pmoA* gene sequences of 25 Operational Taxonomic Units (OTUs), in relation to uncultivated methanotrophic clusters and methanotroph isolates. The tree was built using the LG empirical amino acid substitution model and bootstrapped using 100 bootstrap replicates. Node numbers indicate bootstrap branch support ≥ 60. OTUs retrieved in this study are displayed in bold. Genebank accession numbers for the sequences at individual node tips are given in parentheses. The scale bar displays 0.2 changes per amino acid position.

**Figure 3.**
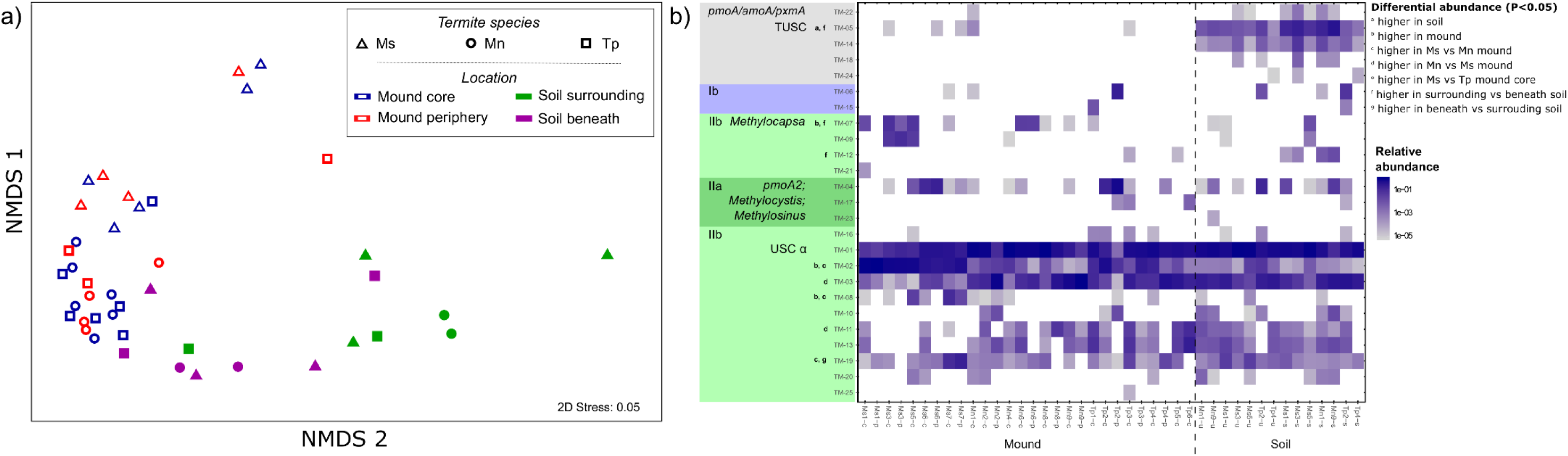
Community composition of methane-oxidising bacteria (methanotrophs) of mounds and adjoining soils. a) The non-metric multidimensional scaling (nMDS) ordination shows the methanotrophic community structure (beta diversity) measured by weighted UniFrac distance metric of the *pmoA* gene. The plot further differentiates the methanotroph communities according to termite species and sample location. b) Heatmap showing the relative abundance of the *pmoA* OTUs in all samples. OTUs are ordered according to their position on the phylogenetic tree shown in Figure 3. Differential abundance of *pmoA* OTUs between sample groups was assessed from negative binomial models of the OTU read counts; *p* values were corrected with the false discovery rate approach to account for multiple testing. Only significant tests are shown.

The 25 OTUs detected were visualised on a phylogenetic tree against reference sequences from a curated *pmoA* gene database [29]. Phylogenetic analysis indicates that all OTUs were affiliated with proteobacterial methanotroph sequences **(Figure 2)**. Across all samples, over 80% of the sequences were affiliated with USCα, a recently cultivated lineage of alphaproteobacterial methanotrophs known to mediate atmospheric CH_4_ oxidation [41, 42] **(Figure 2 & Figure 3b)**. The second most dominant taxonomic groups were affiliated with the alphaproteobacterial lineages *Methylocystis* in mound samples (<10% relative abundance) and the gammaproteobacterial lineage TUSC in soil samples (<10% relative abundance). There was a large proportion of shared taxa across the samples, with the three most abundant OTUs (USCα-affiliated) present in all samples, regardless of type (mound vs soil), location and termite species **(Figure 3b)**. However, differential abundance analysis supported the observed differences between sample type and termite species observed by Unifrac analysis **(Figure 3a)**. Overall, USCα and *Methylocystis* OTUs were more abundant in mound core, mound periphery, and soils beneath, whereas TUSC OTUs were more abundant in surrounding soils. Significant differential abundance was also observed for certain OTUs between termite species **(Figure 3b)**.

It should be noted that community composition of the mounds from the three Australian termite species strikingly differs from those of the African fungus-growing termite *Macrotermes falciger* [14]. These African mounds were dominated by the Jasper Ridge 3 cluster (JR3), a gammaproteobacterial methanotroph lineage closely related to USCγ, which was not detected in the Australian mounds **(Figure 2)**. These differences may reflect the distinct habitat specificity of USCγ and USCα methanotrophs. USCα often occurs in acidic to neutral upland soils ([43], reviewed in [18]), which match the pH values of 5 measured in the mounds of this study **(Table S1)**. In contrast, USCγ and associates lineages are commonly found in upland soils of neutral to basic pH [44, 45], which corresponds well to pH values of 7 to 8 in *Macrotermes falciger* mounds [14].

### Methanotroph communities are kinetically adapted to elevated CH_4_ concentrations

We determined the kinetics of CH_4_ oxidation in the mounds by performing *in situ* GPPTs. Methane oxidation rate was high across the 29 mounds from all three species investigated **(Figure 4a)**. The relationship between CH_4_ concentration and reaction rate best fitted a Michaelis-Menten model for 18 mounds and a linear model for 11 mounds based on AIC values **(Figure 4a)**. For the former group of mounds, apparent Michaelis-Menten coefficients (*K*_m_, *V*_max_) were calculated. Estimated *K*_m_ values for the 18 mounds ranged from 0.32 to 47 µmol (L air)^−1^, and *V*_max_ from 8.4 to 280 µmol (L air)^−1^ h^−1^. These parameters did not significantly differ between termite species. The overall mean values for *K*_m_ and *V*_max_ were 17.5 ± 3.5 µmol (L air)^−1^ and 78.3 ± 17 µmol (L air)^−1^ h^−1^, respectively (standard error of the mean); such values were close to the optimal parameters when fitting a Michaelis-Menten model to combined GPPT data (excluding mounds with linear behavior): *K*_m_ = 13.2 ± 3.5 µmol (L air)^−1^ and *V*_max_ = 55.4 ± 8.5 µmol (L air)^−1^ h^−1^ **(Figure 4a and 4b).** Thus, the methanotroph communities within termite mounds have an apparent medium (µM) affinity for CH_4_. The apparent *K*_m_ is approximately one to two orders of magnitude higher than high-affinity (nM) uptake observed in upland soils [46–48], but one to two orders of magnitude lower than the low-affinity (mM) uptake measured in landfill-cover soils [49]. Similar Michaelis-Menten values were estimated from GPPTs in the vadose zone above a contaminated aquifer (∼1 to 40 µL L^−1^), which featured CH_4_ concentrations in a similar range to termite mounds [35].

**Figure 4.**
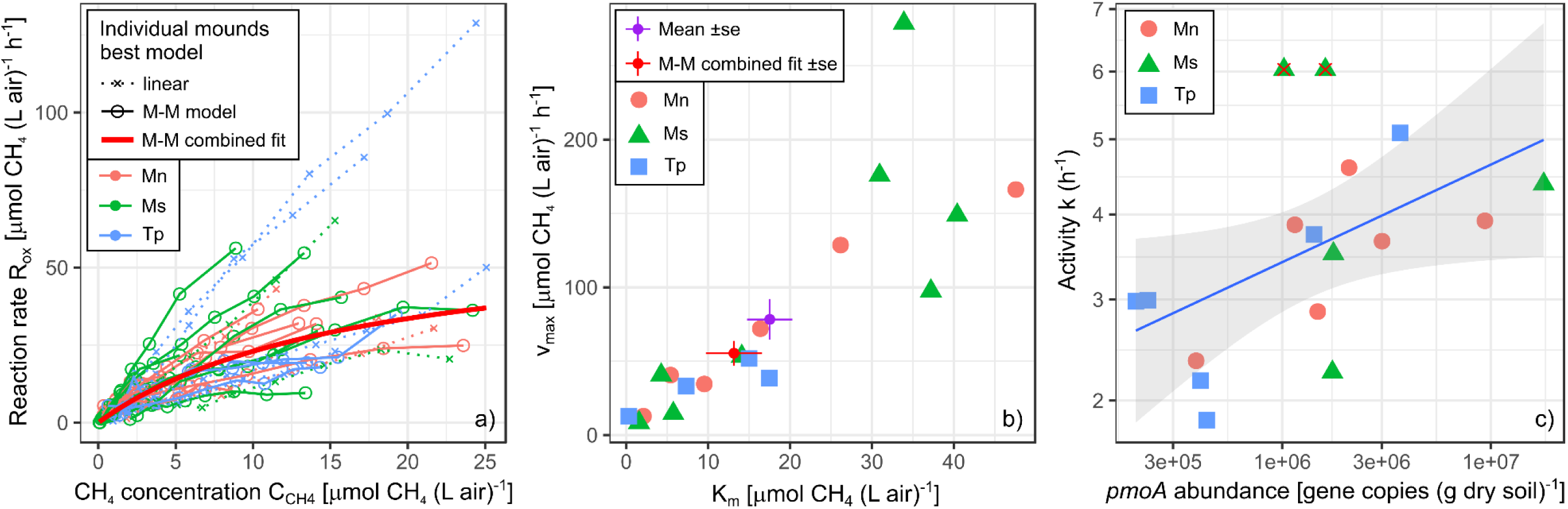
*In situ* kinetic parameters of methane oxidation in termite mounds. a) Optimal Michaelis-Menten (M-M) curve estimated for a combined data set of *in situ* concentrations and reaction rates from 18 out of 29 mounds showing M-M behavior, excluding 11 mounds with an apparent linear increase of rates based on AIC. b) Individual M-M parameters of the 18 mounds showing M-M behavior, including their mean and standard error, and the optimal M-M parameters of the combined data set. c) Positive correlation of activity coefficients *k* with *pmoA* gene abundance in mound core samples of the 17 mounds selected for detailed analysis (R^2^ = 0.44, *p* = 0.0069). Two outliers have been removed, indicated with red X.

It is noteworthy that 11 mounds showed an apparent linear increase of reaction rates with substrate concentrations **(Figure 4a)**. This could indicate that *V*_*max*_ has not been reached during GPPTs with a maximum injected CH_4_ concentration of ∼900 µL L^−1^ (∼40 µM); indeed, the injection concentration is in the range of our highest *K*_*m*_, thus the capacity of some mounds to oxidise CH_4_ can be substantially higher. A linear increase could also indicate a shift in kinetics during the course of the GPPT. It is in the nature of the GPPT that different areas of the mound are exposed to different concentration ranges, depending on their distance to the gas injection/extraction point [34]. Hence, gas extracted at different times may have had a “history” of exposure to methanotroph communities with different kinetics. It is even conceivable that the 1 h long exposure to high injection concentrations around the injection/extraction point triggered the upregulation of a low-affinity methane monooxygenase isozyme, such as those reported in *Methylocystis* sp. SC2 [50].

For the 17 selected mounds investigated in more detail, we analysed how the kinetics and abundance of the methanotroph communities were correlated. While a range of positive correlations were observed, the most robustly supported was the relationship between the first-order activity coefficient *k* and termite mound core *pmoA* copy number (R^2^ = 0.44, *p* = 0.007) **(Figure 4c)**. In contrast, correlations with *pmoA* copy number in periphery and soils were not significant. These observations are in line with previous inferences that mound cores are primarily responsible for CH_4_ oxidation in *M. nervosus* and *M. sunteri* mounds [7]. On this basis, cell-specific CH_4_ reaction rates calculated based on methanotroph abundance of the mound core ranged from 1.8 × 10^−17^ to 1.3 × 10^−14^ mol CH_4_ cell^−1^ h^−1^, one to two orders of magnitude higher than observed in upland soils [38]. Though differences between species were not significant, highest cell-specific rates were calculated for *T. pastinator*, with some values close to a value determined from a landfill-cover biofilter [51]. This would imply that these methanotroph communities operate close to their maximum potential of CH_4_ oxidation. However, it is likely that values for *T. pastinator* are an overestimate given previous studies indicate that most CH_4_ for this species is oxidised in the soil beneath rather than mound itself [7]; thus, for this species but not the two others tested, core *pmoA* numbers underestimate the active methanotroph community involved in mitigating termite CH_4_ emissions.

### Conclusions and perspectives

Overall, our results imply that local environmental concentrations of CH_4_ shape the composition and kinetics of the methanotroph community. Elevated CH_4_ production from termites appears to have selected for a specialised medium-affinity methanotroph community within mounds. The community analysis shows that the methanotroph communities within termite mounds are related to, and likely derived from, those of adjoining soils. However, elevated CH_4_ availability and termite activity have facilitated selection for USCα and *Methylocystis* OTUs, together with exclusion of TUSC OTUs, compared to adjoining soil. These methanotroph communities, with high cell-specific reaction rates and medium affinities for CH_4_, are thus ideally kinetically adapted to grow on termite-derived CH_4_ and in turn reduce atmospheric emissions of this greenhouse gas. Concordant findings were made across three different species, with the strongest relationships between methanotroph abundance and methane oxidation parameters observed for *M. nervosus* and *M. sunteri* mounds.

However, the apparent kinetic parameters of methane oxidation in termite mounds are atypical of the dominant groups of methanotrophs present. It is probable that USCα mediates most methane oxidation in mounds, given their abundance and prevalence in the mound core methanotroph communities, as well as the high cell-specific rates of methane oxidation measured. However, USCα are typically high-affinity methanotrophs associated with the oxidation of CH_4_ at atmospheric concentrations in upland soils. There are several possible explanations for this discrepancy. Firstly, while members of USCα in soil and mound share the same methane monooxygenase, the selective pressure of enhanced methane availability may have caused mound inhabitants to evolve faster-acting, lower-affinity variants. An alternative explanation is community heterogeneity. Any apparent Michaelis-Menten parameter estimates from environmental samples are ‘bulk’ values integrating across the whole microbial community and thus a vast array of potentially different enzymes. Indeed, the observed *in situ* kinetics are also compatible with the coexistence of slow-acting, high-affinity USCα methanotrophs alongside faster-acting, low-affinity other groups. While CH_4_ concentrations in mounds are elevated compared to the atmosphere [7, 52], they are generally within an accessible range for high-affinity methanotrophs (2 – 100 ppmv). *Methylocystis*-like OTUs, as the second most abundant group, are likely to be particularly competitive at high CH_4_ concentrations. Members of this group encode kinetically distinct methane monooxygenase isozymes [50] and have been identified in other soils with elevated methane concentrations [22]. They would therefore have a competitive advantage in termite mounds where CH_4_ availability is elevated and exhibits considerable temporal and spatial variation.

However, despite elevated substrate availability, methanotrophs remain minor members of microbial communities in termite mounds. While they are moderately enriched in mound core than adjoining soil overall, they are actually diminished relative to soil in both the mound core of *T. pastinator* and the periphery of all three species. The observed strong correlation of methanotroph abundance with the porosity of the mound material for all species strengthens the case for habitat porosity as a crucial factor in regulating the methanotrophic community, likely by regulating local CH_4_ availability through diffusion. Another factor driving low methanotroph abundance could be the accumulation of ammonia, which is known to be produced in high levels by termites [53]; it is known that ammonia competitively inhibits methane monooxygenase activity [54, 55] and ammonia levels are major environmental factors regulating methanotroph community in soils [56, 57]. However, while we detected high ammonia concentrations in some mounds **(Table S1)**, this did not correlate with interspecies differences in methanotroph abundance. It is also possible that the methanotrophs are limited by other factors, such as the micronutrients required for methane monooxygenase activity. Ultimately, given methanotrophs remain minor members of the termite mound community, they are only able to mitigate a proportion of the large amounts of methane produced by mound-dwelling termites.

## Acknowledgements

We thank Thanavit Jirapanjawat, Lindsay Hutley and Matthew Northwood for technical and logistical assistance, and Andreas Brune for an inspiring correspondence. This work was supported by Australian Research Council grants (DP120101735 and LP100100073 awarded to S.K.A.), Swiss National Science Foundation Early Postdoc Mobility Fellowships (P2EZP3_178421 awarded to E.C.; P2EZP3_155596 awarded to P.A.N.), an ARC DECRA Fellowship (DE170100310 awarded to C.G.), the Terrestrial Ecosystem Research Network (TERN) OzFlux, and the TERN Australian SuperSite Network.

## Author contributions

P.A.N., E.C., S.K.A., and C.G. designed the study. E.C. and P.A.N. performed field and laboratory work. E.C., P.A.N., and C.G. analyzed data. C.G., E.C., P.A.N., and S.K.A. wrote and edited the paper.

## References

1. Brune A. Methanogenesis in the digestive tracts of insects. Handbook of hydrocarbon and lipid microbiology. 2010. Springer, pp 707–728.

2. Zimmerman PR, Greenberg JP, Wandiga SO, Crutzen PJ. Termites: a potentially large source of atmospheric methane, carbon dioxide, and molecular hydrogen. Science (80-) 1982; 218: 563–565.

3. Rasmussen RA, Khalil MAK. Global production of methane by termites. Nature 1983; 301: 700.

4. Sugimoto A, Inoue T, Tayasu I, Miller L, Takeichi S, Abe T. Methane and hydrogen production in a termite-symbiont system. Ecol Res 1998; 13: 241–257.

5. Brauman A, Kane MD, Labat M, Breznak JA. Genesis of acetate and methane by gut bacteria of nutritionally diverse termites. Science (80-) 1992; 257: 1384– 1387.

6. Kirschke S, Bousquet P, Ciais P, Saunois M, Canadell JG, Dlugokencky EJ, et al. Three decades of global methane sources and sinks. Nat Geosci 2013; 6: 813– 823.

7. Nauer PA, Hutley LB, Arndt SK. Termite mounds mitigate half of termite methane emissions. Proc Natl Acad Sci 2018; 115: 13306–13311.

8. Hanson RS, Hanson TE. Methanotrophic bacteria. Microbiol Rev 1996; 60: 439–471.

9. Reuß J, Rachel R, Kämpfer P, Rabenstein A, Küver J, Dröge S, et al. Isolation of methanotrophic bacteria from termite gut. Microbiol Res 2015; 179: 29–37.

10. Pester M, Tholen A, Friedrich MW, Brune A. Methane oxidation in termite hindguts: absence of evidence and evidence of absence. Appl Environ Microbiol 2007; 73: 2024–2028.

11. Dunfield PF. The Soil Methane Sink. In: Reay D, Hewitt K, Smith K, Grace J (eds). Greenhouse Gas Sinks. 2007. CABI, Wallingford, pp 152–170.

12. Bignell DE, Eggleton P, Nunes L, Thomas KL. Termites as mediators of carbon fluxes in tropical forest: budgets for carbon dioxide and methane emissions. For insects 1997; 109–134.

13. Jamali H, Livesley SJ, Grover SP, Dawes TZ, Hutley LB, Cook GD, et al. The importance of termites to the CH4 balance of a tropical savanna woodland of northern Australia. Ecosystems 2011; 14: 698–709.

14. Ho A, Erens H, Mujinya BB, Boeckx P, Baert G, Schneider B, et al. Termites facilitate methane oxidation and shape the methanotrophic community. Appl Environ Microbiol 2013; 79: 7234–7240.

15. Noirot C, Darlington JPEC. Termite nests: architecture, regulation and defence. Termites: evolution, sociality, symbioses, ecology. 2000. Springer, pp 21–139.

16. Korb J. Termite mound architecture, from function to construction. Biology of termites: a modern synthesis. 2010. Springer, pp 349–373.

17. Jones DT, Eggleton P. Global Biogeography of Termites: A Compilation of Sources. In: Bignell DE, Roisin Y, Lo N (eds). Biology of Termites: A Modern Synthesis. 2011. Springer, Dordrecht, pp 1–576.

18. Knief C. Diversity and habitat preferences of cultivated and uncultivated aerobic methanotrophic bacteria evaluated based on *pmoA* as molecular marker. Front Microbiol 2015; 6: 1346.

19. Nauer PA, Chiri E, Souza D de, Hutley LB, Arndt SK. Rapid image-based field methods improve the quantification of termite mound structures and greenhouse-gas fluxes. Biogeosciences 2018; 15: 3731–3742.

20. Holmes AJ, Costello A, Lidstrom ME, Murrell JC. Evidence that participate methane monooxygenase and ammonia monooxygenase may be evolutionarily related. FEMS Microbiol Lett 1995; 132: 203–208.

21. Costello AM, Lidstrom ME. Molecular characterization of functional and phylogenetic genes from natural populations of methanotrophs in lake sediments. Appl Environ Microbiol 1999; 65: 5066–5074.

22. Henneberger R, Chiri E, Bodelier PEL, Frenzel P, Lüke C, Schroth MH. Field-scale tracking of active methane-oxidizing communities in a landfill cover soil reveals spatial and seasonal variability. Environ Microbiol 2015; 17: 1721–1737.

23. Caporaso JG, Lauber CL, Walters WA, Berg-Lyons D, Lozupone CA, Turnbaugh PJ, et al. Global patterns of 16S rRNA diversity at a depth of millions of sequences per sample. Proc Natl Acad Sci 2011; 108: 4516–4522.

24. Parada AE, Needham DM, Fuhrman JA. Every base matters: assessing small subunit rRNA primers for marine microbiomes with mock communities, time series and global field samples. Environ Microbiol 2016; 18: 1403–1414.

25. Apprill A, McNally S, Parsons R, Weber L. Minor revision to V4 region SSU rRNA 806R gene primer greatly increases detection of SAR11 bacterioplankton. Aquat Microb Ecol 2015; 75: 129–137.

26. Chiri E, Nauer PA, Rainer E-M, Zeyer J, Schroth MH. High temporal and spatial variability of atmospheric-methane oxidation in Alpine glacier-forefield soils. Appl Environ Microbiol 2017; 83: e01139–17.

27. Edgar RC. Search and clustering orders of magnitude faster than BLAST. Bioinformatics 2010; 26: 2460–2461.

28. Wen X, Yang S, Liebner S. Evaluation and update of cutoff values for methanotrophic pmoA gene sequences. Arch Microbiol 2016; 198: 629–636.

29. Dumont MG, Lüke C, Deng Y, Frenzel P. Classification of pmoA amplicon pyrosequences using BLAST and the lowest common ancestor method in MEGAN. Front Microbiol 2014; 5: 34.

30. McMurdie PJ, Holmes S. phyloseq: an R package for reproducible interactive analysis and graphics of microbiome census data. PLoS One 2013; 8: e61217.

31. Urmann K, Gonzalez-Gil G, Schroth MH, Hofer M, Zeyer J. New field method: Gas push− pull test for the in-situ quantification of microbial activities in the vadose zone. Environ Sci Technol 2005; 39: 304–310.

32. Reim A, Lüke C, Krause S, Pratscher J, Frenzel P. One millimetre makes the dierence: high-resolution analysis of methane-oxidizing bacteria and their specific activity at the oxic–anoxic interface in a flooded paddy soil. ISME J 2012; 6: 2128.

33. Raj SS, Sumangala RK, Lal KB, Panicker PK. Gas chromatographic analysis of oxygen and argon at room temperature. J Chromatogr Sci 1996; 34: 465–467.

34. Schroth MH, Istok JD. Models to determine first-order rate coefficients from single-well push-pull tests. Groundwater 2006; 44: 275–283.

35. Urmann K, Schroth MH, Noll M, Gonzalez-Gil G, Zeyer J. Assessment of microbial methane oxidation above a petroleum-contaminated aquifer using a combination of in situ techniques. J Geophys Res Biogeosciences 2008; 113.

36. R Development Core Team. R: A Language and Environment for Statistical Computing. 2017. R Foundation for Statistical Computing, Vienna, Austria.

37. Holt JA. Microbial activity in the mounds of some Australian termites. Appl Soil Ecol 1998; 9: 183–187.

38. Kolb S, Knief C, Dunfield PF, Conrad R. Abundance and activity of uncultured methanotrophic bacteria involved in the consumption of atmospheric methane in two forest soils. Environ Microbiol 2005; 7: 1150–1161.

39. King H, Ocko S, Mahadevan L. Termite mounds harness diurnal temperature oscillations for ventilation. Proc Natl Acad Sci 2015; 112: 11589–11593.

40. Bristow KL, Holt JA. Can termites create local energy sinks to regulate mound temperature? J Therm Biol 1987; 12: 19–21.

41. Tveit AT, Hestnes AG, Robinson SL, Schintlmeister A, Dedysh SN, Jehmlich N, et al. Widespread soil bacterium that oxidizes atmospheric methane. Proc Natl Acad Sci 2019; 201817812.

42. Pratscher J, Vollmers J, Wiegand S, Dumont MG, Kaster A-K. Unravelling the identity, metabolic potential and global biogeography of the atmospheric methane-oxidizing Upland Soil Cluster α. Environ Microbiol 2018; 20: 1016–1029.

43. Knief C, Lipski A, Dunfield PF. Diversity and activity of methanotrophic bacteria in dierent upland soils. Appl Environ Microbiol 2003; 69: 6703–6714.

44. Chiri E, Nauer PA, Henneberger R, Zeyer J, Schroth MH. Soil–methane sink increases with soil age in forefields of Alpine glaciers. Soil Biol Biochem 2015; 84: 83–95.

45. Angel R, Conrad R. In situ measurement of methane fluxes and analysis of transcribed particulate methane monooxygenase in desert soils. Environ Microbiol 2009; 11: 2598–2610.

46. Bender M, Conrad R. Kinetics of CH4 oxidation in oxic soils exposed to ambient air or high CH4 mixing ratios. FEMS Microbiol Lett 1992; 101: 261–270.

47. Nauer PA, Schroth MH. In situ quantification of atmospheric methane oxidation in near-surface soils. Vadose Zo J 2010; 9: 1052–1062.

48. Judd CR, Koyama A, Simmons MP, Brewer P, von Fischer JC. Co-variation in methanotroph community composition and activity in three temperate grassland soils. Soil Biol Biochem 2016; 95: 78–86.

49. Schroth MH, Eugster W, Gómez KE, Gonzalez-Gil G, Niklaus PA, Oester P. Above-and below-ground methane fluxes and methanotrophic activity in a landfill-cover soil. Waste Manag 2012; 32: 879–889.

50. Baani M, Liesack W. Two isozymes of particulate methane monooxygenase with dierent methane oxidation kinetics are found in *Methylocystis* sp. strain SC2. Proc Natl Acad Sci 2008; 105: 10203–10208.

51. Gebert J, Stralis-Pavese N, Alawi M, Bodrossy L. Analysis of methanotrophic communities in landfill biofilters using diagnostic microarray. Environ Microbiol 2008; 10: 1175–1188.

52. Jamali H, Livesley SJ, Hutley LB, Fest B, Arndt SK. The relationships between termite mound CH4/CO2 emissions and internal concentration ratios are species specific. Biogeosciences 2013; 10: 2229–2240.

53. Rong JI, Brune A. Nitrogen mineralization, ammonia accumulation, and emission of gaseous NH3 by soil-feeding termites. Biogeochemistry 2006; 78: 267–283.

54. Schnell S, King GM. Mechanistic analysis of ammonium inhibition of atmospheric methane consumption in forest soils. Appl Environ Microbiol 1994; 60: 3514– 3521.

55. Carlsen HN, Joergensen L, Degn H. Inhibition by ammonia of methane utilization in *Methylococcus capsulatus* (Bath). Appl Microbiol Biotechnol 1991; 35: 124– 127.

56. Bodelier PLE, Laanbroek HJ. Nitrogen as a regulatory factor of methane oxidation in soils and sediments. Fems Microbiol Ecol 2004; 47: 265–277.

57. Veraart AJ, Steenbergh AK, Ho A, Kim SY, Bodelier PLE. Beyond nitrogen: The importance of phosphorus for CH<inf>4</inf> oxidation in soils and sediments. Geoderma 2015; 259–260.

