## Supplementary material for "Termite mounds contain distinct methanotroph communities that are kinetically adapted to elevated methane concentrations"

### Supplemental material

**Table S1.** Soil physicochemical data for termite mound core, termite mound periphery, and soil samples. A composite sample was analysed for each species and location.

| | pH ($H_2O$) | TOC | TC | TN | $NH_4^+$-N | $NO_3^-$-N | $PO_4^{2-}$ | $K^+$ |
| --- | --- | --- | --- | --- | --- | --- | --- | --- |
| | | % (kg $kg^{-1}$) | | | mg $kg^{-1}$ | | | |
| Tp-p | 5.4 | 1.86 | 2.05 | 0.09 | 16 | 6 | 6 | 99 |
| Tp-c | 5.4 | 2.11 | 2.36 | 0.10 | 15 | 6 | 5 | 106 |
| Mn-p | 5.5 | 5.04 | 7.15 | 0.25 | 32 | 9 | 9 | 187 |
| Mn-c | 5.8 | 4.83 | 6.68 | 0.24 | 39 | 3 | 8 | 206 |
| Ms-p | 5.2 | 5.05 | 9.96 | 0.42 | 280 | 80 | 40 | 140 |
| Ms-c | 5.2 | 4.41 | 10.7 | 0.44 | 220 | 36 | 29 | 128 |
| Soil | 5.7 | 2.38 | 3.25 | 0.20 | 22 | 7 | 10 | 50 |

**Figure S1.** Summary of differences in the abundance and richness of methanotrophs within different termite mound and adjoining soil samples. a) Abundance of the methanotroph community, based on *pmoA* gene copy number. b) Abundance of the total bacterial and archaeal community, based on 16S rRNA gene copy number. c) Estimated richness of the methanotroph community, based on Chao1 index of the *pmoA* gene. Mound samples (core and periphery) and adjoining soil samples (surrounding and beneath the mound) were tested from mounds three different termite species. Different letters denote significant differences in abundance or richness between sample groups ($p < 0.05$, Wilcoxon signed-rank test).

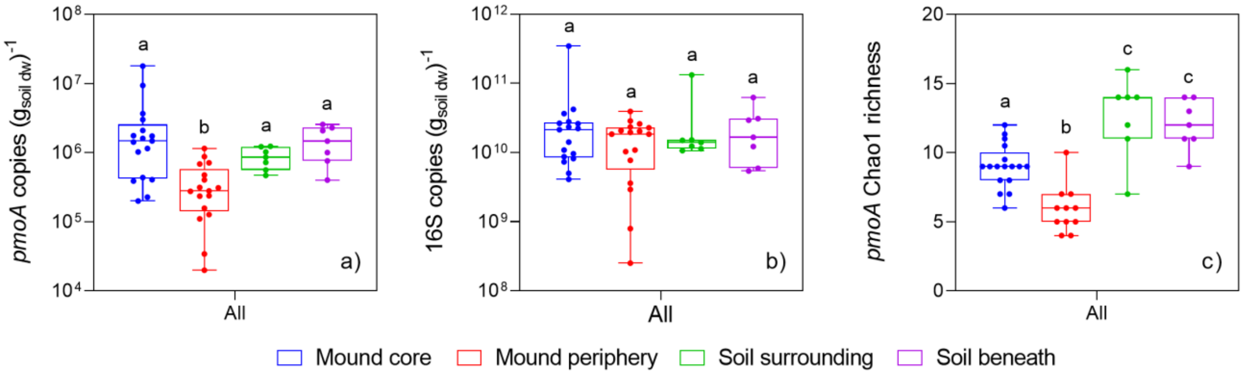

**Figure S2.** Summary of differences in the abundance and richness of methanotrophs within termite mound samples between different termite species. a) Abundance of the methanotroph community, based on *pmoA* gene copy number. b) Abundance of the total bacterial and archaeal community, based on 16S rRNA gene copy number. c) Estimated richness of the methanotroph community, based on Chao1 index of the *pmoA* gene. Mound samples (core and periphery) were tested from mounds three different termite species, *Microcerotermes nervosus* (Mn), *Macrognathotermes sunteri* (Ms), and *Tumulitermes pastinator* (Tp). Different letters denote significant differences in abundance or richness between sample groups ($p < 0.05$, Wilcoxon signed-rank test).

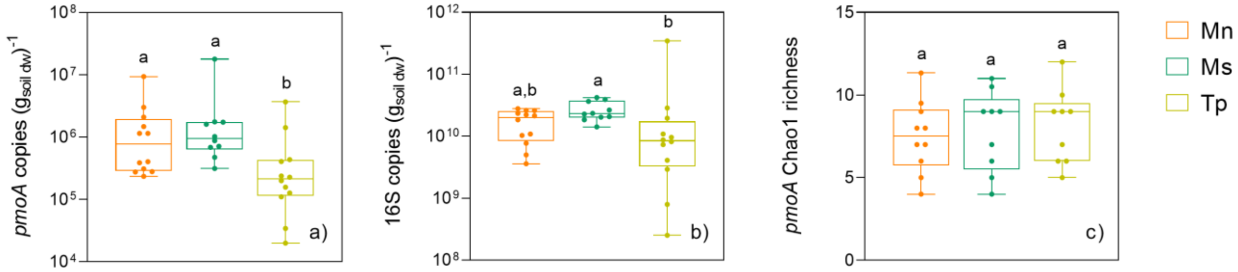

**Figure S3.** Summary of differences in the diversity of methanotrophs within different termite mound and adjoining soil samples. a) Shannon diversity of the *pmoA* gene. b) Inverse Simpson diversity of the *pmoA* gene. Mound samples (core and periphery) and adjoining soil samples (surrounding and underlying mound) were tested from mounds three different termite species. Different letters denote significant differences in abundance or richness between sample groups ($p < 0.05$, Wilcoxon signed-rank test).

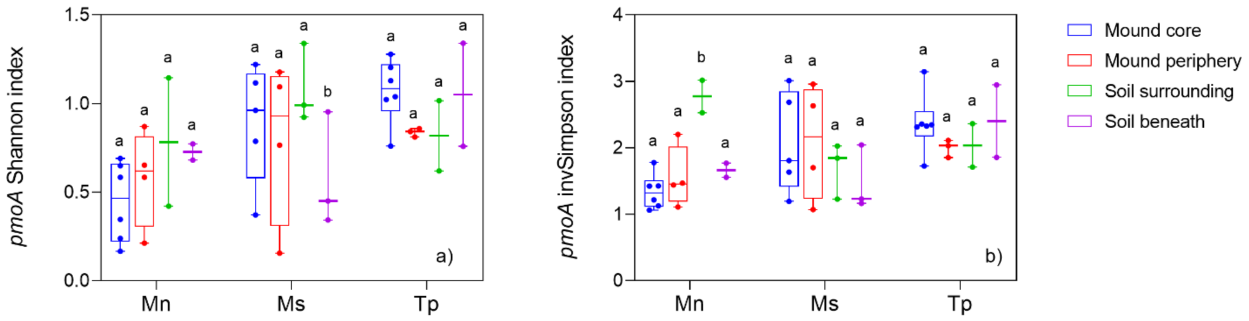
